# Biomarker-based assessment of the muscle maintenance and energy status of anurans from an extremely seasonal semi-arid environment, the Brazillian Caatinga

**DOI:** 10.1101/394544

**Authors:** Carla Bonetti Madelaire, Fernando Ribeiro Gomes, Inna Sokolova

## Abstract

Strongly seasonal environments pose challenges for performance and survival of animals, especially when resource abundance seasonally fluctuates. We investigated the seasonal variation of expression of key metabolic biomarkers in the muscles of three species of anurans from the drastically seasonal Brazilian semi-arid area, Caatinga. The three studied anuran species (*Rhinella jimi, R. granulosa* and *Pleurodema diplolister*) differ in their seasonal activity patterns. We examined the expression of proteins regulating energy turnover (AMP-activated protein kinase [AMPK] and protein kinase B [AKT]), protein synthesis and homeostasis (total and phosphorylated eukaryotic initiation factor 2α [eIF2α and p-eIF2α] and chaperone proteins [HSP 60, 70, and 90]) in muscles related to reproduction and locomotion. Cytochrome c oxidase (COX) activity was also assessed as an index of the muscle aerobic capacity. Our results point to the importance of metabolic regulators mediating the muscular function during the drastic seasonal variation. The toads that remain active during the drought appear to maintain muscles through more energy extensive pathways including elevated protein synthesis, while the aestivating species employs energy conservation strategy suppressing protein synthesis, decreasing chaperone expression and increasing expression of AMPK. All three studied species activate cell survival pathways during the drought likely to prevent muscle atrophy, and maintain the muscle capacity throughout the year, despite the resource limitation. These strategies are important considering the unpredictability of the reproductive event and high demand on muscular activity during the reproductive season in these amphibians.

**Summary statement:** We studied seasonal variation of key metabolic regulators in the muscles of anurans that experience drastic variation in environmental conditions and differ in the seasonal activity patterns.

## Introduction

Strongly seasonal environments characterized by large changes in conditions such as temperature and food availability, pose challenges to the organisms that need to adjust their functions to survive and complete their life cycles [1, 2]. These physiological adjustments are dependent on biochemical regulators and cell signaling pathways that mediate cell survival, ensure cell and organ integrity and prioritize the use of energy to meet the (often conflicting) needs of survival, growth and reproduction [1,3]. In vertebrate ectotherms, the skeletal muscle physiology and performance are directly related to the overall fitness due to the muscles’ involvement in courtship, territorial defense, foraging, escape from predators, mating interactions and migration [4]. In seasonally fluctuating environments, skeletal muscles can display phenotypic changes such as atrophy and changes in the fiber number and type during extreme environmental stress and/or resource limitation [5]. Nevertheless, anuran species that aestivate display little or no changes in muscle morphology and performance [6,7] suggesting that species living in arid and semi-arid environments may have molecular regulatory mechanisms supporting the muscle maintenance and functionality under the unfavorable environmental conditions.

There is an extensive literature on physiological and biochemical adjustments associated with metabolic depression in anurans from environments with strongly seasonal and/or unpredictable climatic variation [8,1,2]. However, possible biochemical adjustments displayed by anurans that remain active during the periods of severe environmental stress and resource limitation are not well understood. Furthermore, different locomotor muscles can display distinct reduction in cross-sectional area and mechanical proprieties in aestivating anuran species and mammalian hibernators, prioritizing the maintenance of the muscles most relevant to performance [7,9]. Yet, it is unknown whether seasonal adjustments differ between reproductive and locomotor muscle in anurans, reflecting preferential investment into the maintenance and reproductive effort. The possible molecular adjustments of locomotor and reproductive muscles that help support individual’s performance and survival [1] under environmental stress and resource limitation are not well understood and require further investigation.

Anurans from the Brazilian semi-arid area, the Caatinga, face drastic seasonal changes in environmental conditions [10-12]. In the Caatinga, anurans depend on unpredictable heavy rain events (during the short rain season between January and April) to reproduce. Occasional small showers during the rainy season are not sufficient to trigger breeding, yet make water and food more easily available to anurans. The rest of the year is the dry season, characterized by scarce water and food resources. In some years, there is no rain so that the anurans have no reproductive opportunity until next rainy season. Anurans from the Caatinga show interspecific variation in behavioral strategies to survive the drought. Some species such as *Rhinella jimi* and *R. granulosa* remain active and foraging around humid areas [13]. In contrast, *Pleurodema diplolister* aestivates and does not feed until the first rains start [12; 14]. Previous studies show that these anuran species display adjustments in reproductive [13] and immune physiology [14] throughout the year. During the reproductive period, they display elevated plasma levels of androgens and higher immunological profile and response compared with the dry season [14]. During the drought, anurans present lower steroid plasma levels and lower immune parameters and performance, with the strongest suppression observed in the aestivating species [13,14]. This stark seasonality of the physiology and behavior in the anurans from the semi-arid Caatinga suggests a strong selective pressure to regulate energy metabolism and muscle function to meet the seasonally variable demands of reproduction, resource acquisition and activity (including the physiological challenge of metabolic depression during aestivation).

To assess the potential mechanisms involved in the regulation and maintenance of the muscle function in anurans from an extremely seasonal environment of Brazilian semi-arid Caatinga, we investigated the seasonal variation of expression of key metabolic regulators in the muscles of males from three species of anurans that differ in their seasonal activity patterns: *Rhinella jimi* and *R. granulosa* that remain foraging during drought and *P. diplolister,* which aestivates during this period. We studied muscles predominantly specialized on reproduction (including the trunk and larynx muscles used to sustain calling activity [15,16] and the flexor carpi radialis used in amplexus (grasping) behavior [17]) as well as the muscles predominantly specialized on locomotion (the plantaris muscle [18]). During dry season, we expected to see an increase in the pathways that support cell survival and stress response in the muscle, along with the suppression of the protein synthesis and catabolic pathways [3], and a decrease of muscle aerobic capacity to conserve energy [19], especially in muscles related to reproduction. The prioritization of the muscles integrity during drought might ensure the rapid mobilization of these muscles during an unpredictable reproductive opportunity, and thus would be adaptive. We also anticipated that the aestivating species would display a more intense downregulation of metabolic functions, such protein synthesis, compared with the species that remain active year round.

To test these hypotheses, we investigated the expression of two protein kinases that play a key role in the regulation of energy turnover in the muscle (the AMP-activated protein kinase and protein kinase B) and key proteins involved in the regulation of the protein synthesis and homeostasis (eukaryotic initiation factor 2α and chaperone proteins). The cytochrome c oxidase (COX) activity was assessed as an index of the mitochondrial aerobic capacity in the tissue [20]. The AMP-activated protein kinase (AMPK) is an energy sensor of the cell responding to the AMP:ATP ratio [21,22]. During periods that animals need to save energy, AMPK is activated to regulate catabolic *versus* anabolic metabolism increasing ATP synthesis and suppressing ATP consumption [21,22]. Protein kinase B (AKT) plays a central role in metabolism and cell survival stimulating glucose uptake, glycogen synthesis, lipogenesis and protein synthesis, and regulating the cell cycle and apoptosis [23]. Eukaryotic initiation factor 2α regulates protein synthesis and plays a key role in the stress response and suppression of the ATP-consuming protein synthesis under low energetic budget scenarios [24]. Heat shock proteins are involved in the general stress response acting as molecular chaperones and regulating folding of newly synthesized proteins or those damaged by stressors [25]. Considering their key roles in regulation of the muscle integrity and function, we expect to see different patterns of activation of the studied signaling proteins across the season in different species. We also anticipate that the stress-related pathways involved in energy conservation are upregulated and the aerobic capacity (measured as the COX activity) is suppressed in anuran muscles during the drought, when the species that remain active (*R. jimi* and *R. granulosa*) are facing food and water shortage, and *P. diplolister* is aestivating.

## Materials and Methods

### Field collections

Field work was conducted at Fazenda São Miguel near the city of Angicos, in the State of Rio Grande do Norte, Brazil (5°30’43”S. 36°36’18’’W). The area is in the domain of Brazilian Caatinga, and is characterized by high temperatures. January is the hottest month with an average temperature of 27.4°C (minimum: 22.8°C, maximum: 32.0°C), and July is the coldest month, with an average temperature of 24.3°C (minimum: 20.3°C, maximum: 28.3°C) (http://pt.climate-data.org/location/312354/). The annual average temperature from 1950 to 2000 is 26.6 °C (worldclim.org) and there are two distinct seasons: a rainy season (January to April, 96.4 mm of precipitation/month) and a dry season (August to November, 2.5 mm of precipitation/month). The intermediate months of June, July and December can be considered dry or rainy depending on the extent of drought in the specific year. In response to the challenges of the dry season, anurans from this locality have adopted different behavioral strategies. *R. granulosa* and *R. jimi* remain active, foraging close to humid areas and artificial water sources [14], while *Pleurodema diplolister* aestivate in the sandy soil under the beds of temporary rivers [12]. For this study, animals were collected during two different periods in 2015: (A) during the reproductive season (March, 5–12^th^, 2015); (B) during the dry season (August, 10–16^th^, 2015).

During the reproductive period, males of *R. granulosa* (N = 13), *R. jimi* (N = 11) and *P. diplolister* (N = 16) were located by visual inspection. During the dry period, males of *R. granulosa* (N = 6), *R. jimi* (N = 7) were found by visual inspection, and *P. diplolister* (N = 8) was found by excavating the known burrowing sites in the sandy soil. Anurans were collected and individually maintained in plastic containers with access to water, except the individuals of *P. diplolister* collected during the dry period, which were maintained in plastic containers filled with humid sand collected in the location they were found. After two days, animals were weighted (to the nearest 0.01g) and euthanized with an injection of sodium thiopental solution (25 mg/ml) (Thiopenthax) while kept in an ice-cold dry bath. The muscles (plantaris from the posterior limb, flexor from anterior limbs, larynx and trunk) were rapidly dissected. Muscle samples were either immediately frozen in liquid nitrogen (for immunoblotting analyses), or incubated for 1 to 2 minutes in a cryopreservation medium (10 mM EGTA, 1.3 mM CaCl_2_, 20 mM imidazole, 20 mM taurine, 49 mM K-MES, 3 mM K_2_HPO_4_, 9.5 mM MgCl_2_, 5 mM ATP, 15 mM phosphocreatine, 10 mg/ml fatty acid-free BSA, 20% glycerol, pH 7.1) [26] prior to freezing (for COX activity). Tissues were stored in liquid nitrogen until their transport to the University of North Carolina at Charlotte (UNC Charlotte), NC, USA on dry ice. At UNC Charlotte the muscle samples were stored at ‒80°C until analyses. Fieldwork, maintenance of animals, and transport of samples were conducted under the approved permissions of Comissão de Ética no Uso de Animais do IB (CEUA) (Protocol number: 181/2013) and Ministério do Meio Ambiente, ICMBio, SISBio (License to collect and transport animals: N°29896-1; Export License number: 15BR017888/DF).

### Immunoblotting

Muscle samples were homogenized (1:10 w:v) in ice-cold buffer (100 mM Tris, pH = 7.4, 100 mM NaCl, 1 mM EDTA, 1 mM EGTA, 1% Triton-X, 10% glycerol, 0.1% sodium dodecylsulfate [SDS], 0.5% deoxycholate, 0.5 μg mL^−1^ leupeptin, 0.7 μg mL^−1^ pepstatin, 40 μg mL^−1^ phenylmethylsulfonyl fluoride [PMSF], and 0.5 μg mL^−1^ aprotinin), sonicated three times for 10 s each (output 69 Watts; Sonicator 3000, Misonix Inc.) and centrifuged at 14000 × g for 5 min at 4°C. The protein content of the supernatant was measured using Bio-Rad Protein Assay kit (Bio-Rad, Hercules, CA, USA) with the bovine serum albumin (BSA) as a standard. Protein-containing supernatant was mixed 3:1 (v:v) with a solution containing 4 parts of 4x Laemmli buffer and 1 part of 1 M dithiothreitol (DTT), boiled for 5 minutes and frozen in ‒20°C until further analysis.

Samples (20-50 µg protein per lane, depending on the antibody) were loaded into 10% polyacrylamide gels and run at 72V for 3 hours at room temperature. After the run, the gels were incubated for 30 min in 96·mmol·l ^−1^ glycine, 12·mmol·l ^−1^ Tris and 20% methanol (v/v). The proteins were transferred to a nitrocellulose (for HSP60, HSP70 and HSP90) or polyvinylidene difluoride (PVDF) membrane (for all other antibodies) using a Trans-Blot semi-dry cell (Thermo Fisher Scientific Inc., Portsmouth, NH, USA). Membranes were blocked for one hour in 3% non-fat milk in Tris-buffered saline, pH 7.6 with 0.1% Tween 20 (TBST) at room temperature, and incubated overnight at 4°C with the primary antibodies diluted 1:1000 in 5% BSA in TBST. After washing off the primary antibody with TBST, the membranes were probed with the polyclonal secondary antibodies conjugated with horseradish peroxidase (Jackson ImmunoResearch, West Grove, PA, USA) diluted 1:1000 with 3% non-fat milk in TBST for one hour at the room temperature. After washing off the secondary antibody, the proteins were detected using enhanced chemiluminescence according to the manufacturer’s instructions (Amersham Biosciences, Pierce, Rockford, IL, USA). The signals were captured on X-ray film and relative optical density of protein bands was digitalized with an image analysis software (Gel Doc EZ Imager, Bio-Rad, Hercules, CA, USA) and quantified using Image Lab^™^ software (Bio-Rad Laboratories Inc., Hercules, CA, USA). The loading order of samples in the gels was randomized. The protein loads per lane were identical for all muscle types and for both studied seasons except for p-eIF2α in *R. granulosa* collected during the dry season where the original load of 30 µg per lane did not produce a signal, and 50 µg per lane was used. A single sample (used as an internal control) was loaded on each gel and used to standardize the expression of the target proteins and account for the potential gel-to-gel signal variation. The following antibodies were used: AKT (AKT rabbit polyclonal IgG; Cell Signalling Technology, cat. #9272, Danvers, MA, USA), total AMP-activated protein kinase (AMPK) (AMPKα, Thr172, Rabbit mAb, Cell Signalling Technology, cat. #2535, Danvers, MA, USA), phospho-EIF-2α (Ser51) (no. 07-760, Millipore, cat. #07-760, Temecula, CA, USA), EIF-2α (no. AHO1182, Life Technology, Grand Island, NY, USA), HSP 60 (HSP60 [insect] polyclonal antibody, Enzo Life Science, cat.#ADI-SPA-805-D, Farmingdale, NY, USA), HSP 70 (Heat Shock Protein 70 [HSP70] Ab-2, Mouse Monoclonal Antibody, Thermo Fisher Scientific Inc., cat. # MA3-006, Portsmouth, NH, USA), HSP 90 (Anti-HSP90 antibody, Rat [monoclonal], cat. #SPA-835, Stressgen Bioreagents, Ann Arbor, MI, USA). All antibodies produced a single band of the expected length (S1 Fig).

### Measurements of cytochrome c oxidase capacity

Cryotubes containing the frozen plantaris or trunk muscles were incubated for ~ 2 min at 35°C until the cryopreservation medium was completely thawed. The muscle fibers were immediately washed in ice-cold medium containing 120 mM KCl, 10 mM NaCl, 2 mM MgCl_2_, 2 mM KH_2_PO_4_, 20 mM HEPES, 1mM EGTA Ca-free, 10µg mL^−1^ PMSF and homogenized in 1-2 ml of the same media with several passes of a Potter-Elvenhjem homogenizer and a loosely fitting Teflon pestle at 200 rpm. The homogenate was centrifuged at 2000 × g and 4°C for 8 min to remove cell debris, and the supernatant containing mitochondria was used to measure activity of cytochrome c oxidase (COX).

COX activity was determined by measuring the oxygen consumption of mitochondria-containing supernatant at 23°C in the presence of 5 µM antimycin A, 5 mM ascorbate and 10 mM *N,N,N*′,*N*′-Tetramethyl-*p*-phenylenediamine (TMPD) as an electron donor [27]. Oxygen concentrations were monitored using a fiber optic oxygen sensor connected to the Microx TX3 oxygen monitor with temperature correction (Precision Sensing, Dussel-dorf, Germany) and Oxy Micro ver. 2.00 software (World Precision Instruments, Sarasota, FL). A two-point calibration was performed prior to each measurement. To correct for the potential autooxidation of TMPD, oxygen consumption was measured after addition of 25 mM KCN to inhibit COX, and the difference in the oxygen consumption rates in the presence and absence of KCN was used to calculate COX activity. Concentrations of oxygen in the respiration chamber were monitored using Logger Pro 3.2 with a Vernier LabPro interface (Vernier Software and Techi8nology, Beaverton, OR). Protein concentrations in mitochondrial isolates were measured using a Bio-Rad protein assay (Bio-Rad, Hercules, CA, USA) in the presence of 0.1% Triton X-100 to solubilize mitochondrial membranes, with BSA as the standard. COX activity was expressed as µmol O_2_ min^−1^ g^−1^ mitochondrial protein. Each biological replicate represented an individual isolate obtained from the pooled tissues of 2–3 animals.

### Statistical analysis

Variables were log_10_ transformed to improve normality. For the immunoblotting results, a two-way ANOVA was used to compare the effects of the season, gel and their interaction on protein expression (measured as a densitometric signal). When the effects of the gel and/or gel x season interactions were not significant, a one-way ANOVA was used to test for the effect of the season. To compare different muscles within a season, a one-way ANOVA was used followed by Bonferroni post-hoc test. For COX activity, a two-way ANOVA was used to compare the effects of species and muscles (COX ~ species + muscle); and species and season in each muscle (COX_muscle_ ~ species + season). Assumptions of the ANOVA and t test (normality and equal variance) were met and statistical significance was set at P < 0.05. All tests were run in R software, version 2.10.0 (R Development Core Team, 2010). Data are shown as means ± the standard error of the mean (S.E.M).

## Results

### Metabolic signaling

Total AMPK showed higher expression during the dry period compared to the reproductive period in all studied muscle types of *R. jimi* (larynx - F_1,6_ = 0.25, P = 0.001; trunk - F_1,12_ = 0.68, P = 0.001; flexor - F_1,9_ = 0.13, P = 0.04; plantaris - F_1,12_ = 0.65, P = 0.003) (Fig 1). A similar trend was seen in *P. diplolister,* albeit it was only significant in the plantaris muscle (F_1,10_ = 0.14, P = 0.002) (Fig 1C). *R. granulosa* displayed similar AMPK expression levels across the two studied seasons in all muscle types (Flexor: F_1,8_ = 2.74, P = 0.14; trunk: F_1,12_ = 0.84, P = 0.38; plantaris: F_1,12_ = 0.34 P = 0.57). When compared within the same season, AMPK levels in different muscle types were similar within *R. jimi* and *R. granulosa* (P > 0.05, Table 1). In *P. diplolister,* AMPK levels were similar in different muscle types during the dry period (Table 1). During the reproductive period, trunk muscles of *P. diplolister* showed higher AMPK expression compared with other muscles (F_2,20_ = 4.23, P = 0.03, Bonferroni P = 0.03) (S2 Fig).

**Table 1.** ANOVA: comparison of protein expression in different muscles within a season. F ratios (with the degrees of freedom and the error shown as a subscript), P values are given, and significant effects (P < 0.05) are highlighted in bold. Missing analysis are either protein detection below limit or lack of samples.

|  | *Rhinella jimi* | *Rhinella granulosa* | *Pleurodema diplolister* |
| --- | --- | --- | --- |
|  | Reproductive | Reproductive | Reproductive |
| AKT | **$F_{3,13} = 3.69$ $P = 0.04$** | $F_{3,13} = 3.35$ $P = 0.053$ | $F_{3,11} = 0.91$ $P = 0.47$ |
|  | Dry | Dry | Dry |
| | $F_{3,6} = 4.56$ $P = 0.06$ | $F_{3,5} = 0.34$ $P = 0.80$ | $F_{3,10} = 1.39$ $P = 0.30$ |
| AMPK | Reproductive | Reproductive | Reproductive |
| | $F_{3,18} = 2.44$ $P = 0.10$ | $F_{3,13} = 1.70$ $P = 0.20$ | $F_{3,17} = 3.42$ $P = 0.04$ |
|  | **Dry** | **Dry** | **Dry** |
| | $F_{3,18} = 2.35$ $P = 0.11$ | $F_{2,7} = 0.72$ $P = 0.52$ | $F_{1,10} = 0.14$, $P = 0.002$ |
|  | **Reproductive** | **Reproductive** | **Reproductive** |
| **p-eIF2α** | $F_{3,14} = 2.21$ $P = 0.13$ | $F_{3,6} = 0.87$ $P = 0.51$ | $F_{3,15} = 1.59$ $P = 0.24$ |
|  | **Dry** | **Dry** | **Dry** |
| | $F_{3,14} = 0.74$ $P = 0.55$ | - | $F_{3,6} = 11.53$ $P = 0.01$ |
|  | **Reproductive** | **Reproductive** | **Reproductive** |
| **eIF2α** | $F_{3,15} = 11.18$ $P = 0.001$ | $F_{3,18} = 23.82$ $P = 0.001$ | $F_{3,19} = 2.89$ $P = 0.06$ |
|  | **Dry** | **Dry** | **Dry** |
| | $F_{3,5} = 26.67$ $P = 0.002$ | $F_{3,6} = 1.19$ $P = 0.39$ | $F_{3,5} = 1.02$ $P = 0.46$ |
|  | **Reproductive** | **Reproductive** | **Reproductive** |
| **p-eIF2α/eIF2α** | $F_{3,10} = 1.94$ $P = 0.19$ | $F_{3,5} = 0.17$ $P = 0.92$ | $F_{3,12} = 1.28$ $P = 0.32$ |
|  | **Dry** | **Dry** | **Dry** |
| | $F_{3,6} = 11.30$ $P = 0.007$ | - | $F_{3,4} = 3.02$ $P = 0.16$ |
|  | **Reproductive** | **Reproductive** | **Reproductive** |
| **HSP60** | $F_{3,22} = 9.44$ $P = 0.001$ | $F_{3,15} = 3.70$ $P = 0.04$ | $F_{2,13} = 1.46$ $P = 0.27$ |
|  | **Dry** | **Dry** | **Dry** |
| | $F_{3,12} = 13.90$ $P = 0.001$ | $F_{2,6} = 0.002$ $P = 0.99$ | $F_{3,5} = 2.65$ $P = 0.16$ |
|  | **Reproductive** | **Reproductive** | **Reproductive** |
| **HSP70** | $F_{3,16} = 10.36$ $P = 0.001$ | $F_{3,9} = 0.47$ $P = 0.71$ | $F_{2,25} = 0.92$ $P = 0.41$ |
|  | **Dry** | **Dry** | **Dry** |
| | $F_{3,6} = 0.90$ $P = 0.50$ | $F_{3,8} = 1.16$ $P = 0.38$ | $F_{3,8} = 2.63$ $P = 0.12$ |
|  | **Reproductive** | **Reproductive** | **Reproductive** |
| **HSP90** | $F_{3,12} = 28.07$ $P = 0.001$ | - | $F_{2,8} = 40.87$ $P = 0.001$ |
|  | **Dry** | **Dry** | **Dry** |
| | $F_{3,14} = 1.45$ $P = 0.27$ | - | $F_{1,4} = 0.003$ $P = 0.96$ |

**Fig 1.**
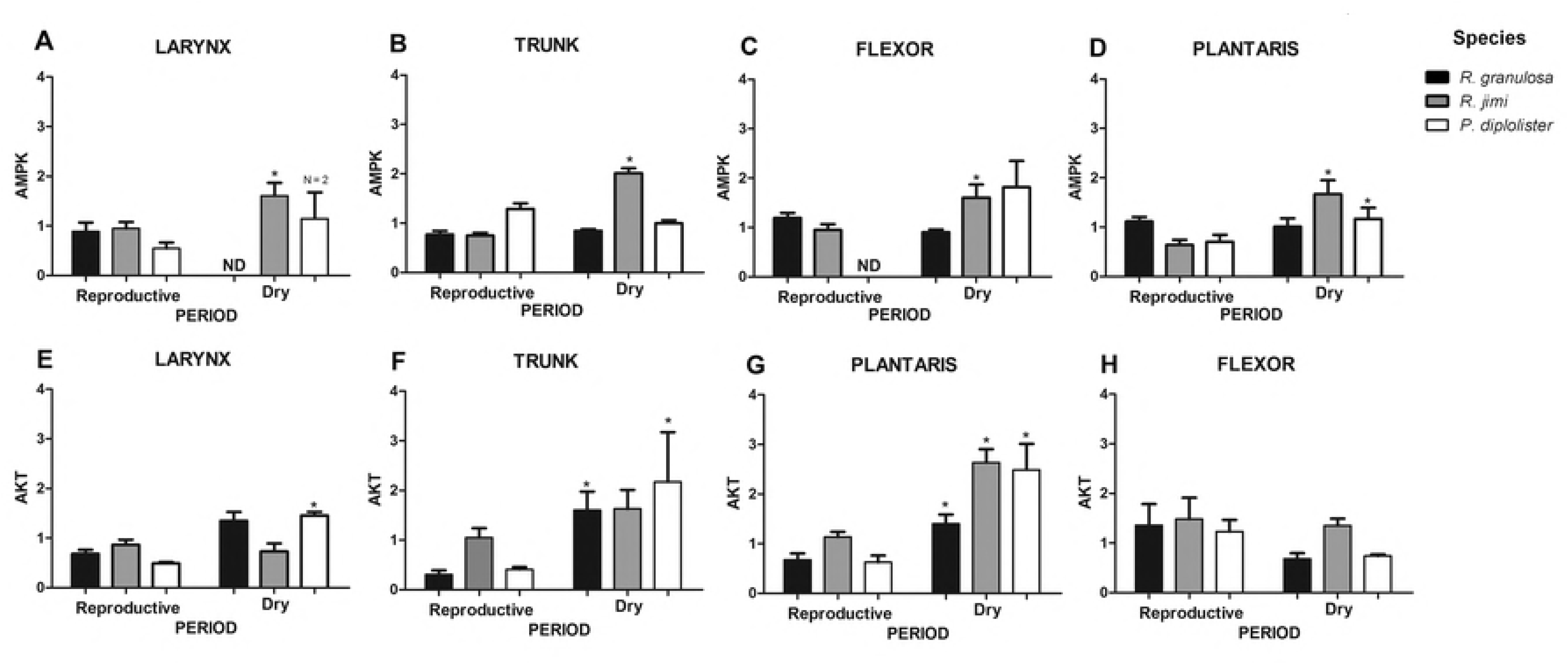
Expression of AMPK and AKT in different muscle types of *R. granulosa, R. jimi*, and *P. diplolister* during reproductive and dry periods. n = 3–10. AMPK expression is shown for (A) larynx, (B) trunk, (C) plantaris, and (D) flexor muscles. AKT expression is shown for (E) larynx, (F) trunk, (G) plantaris, and (H) flexor muscles. Asterisks indicate significant difference between the reproductive and dry periods (P < 0.05). ND - not determined because of the lack of samples.

Expression of the protein kinase B (AKT) tended to be higher during the dry period in all studied species and muscle types (except flexor for all species and the larynx of *R. jimi;* Fig 1E-H). This trend was significant in the larynx muscles from *P. diplolister* (F_1,8_ = 0.58, P = 0.001), the trunk muscles from *R. granulosa* and *P. diplolister* (F_1,7_ = 1.55, P = 0.002; F_1,6_ = 0.79, P = 0.006), and the plantaris muscle from all three studied species (*R. granulosa* - F_1,7_ = 0.24, P = 0.03; *R. jimi* - F_1,6_ = 0.25, P = 0.001; *P. diplolister* - F_1,11_ = 1.21, P = 0.002) (Fig 1E-G). There were no significant differences of AKT expression between seasons in the flexor muscle in the three studied species (*R. granulosa* - F_1,6_ = 3.57, P = 0.11; *R. jimi* - F_1,8_ = 0.19, P = 0.68; *P. diplolister* - F_1,4_ = 5.58, P = 0.08). AKT levels were the same across different muscle types when compared within the same season in all three studied species (P > 0.05, Table 1).

### Protein homeostasis

Total expression levels of the eukaryotic initiation factor 2α (eIF2α) were lower during the dry period than in the reproductive period in the larynx and flexor muscles from *P. diplolister* (F _1,7_ = 0.079, P = 0.016; F _1,4_ = 0.110, P = 0.04) (Fig 2A-D), and in the trunk and plantaris muscles from *R. granulosa* (F_1,14_ = 1.06, P = 0.002; F_1,12_ = 0.132, P = 0.02) (Fig 2A-D). In all other muscle types (including the trunk and plantaris of *P. diplolister,* the larynx and flexor of *R. granulosa*, and all muscle types of *R. jimi*), season had no significant effect on eIF2α expression (P > 0.05). In *R. granulosa*, during the dry period, the flexor muscle had the highest levels of total eIF2α when compare with the other studied muscles types; during reproductive period, the flexor and trunk muscles had elevated levels of eIF2α compared larynx and plantaris (Table 1; S3A and B Fig; Bonferroni P = 0.001 and 0.028, respectively). For *R. jimi* collected in the reproductive period, trunk and plantaris muscles showed lower total eIF2α expression compared with the larynx and flexor (Table 1; S3C Fig; Bonferroni P = 0.042). No differences in total eIF2α levels were found between different muscle types of *R. jimi* during the dry period, or of *P. diplolister* during the dry and reproductive periods (Table 1).

**Fig. 2.**
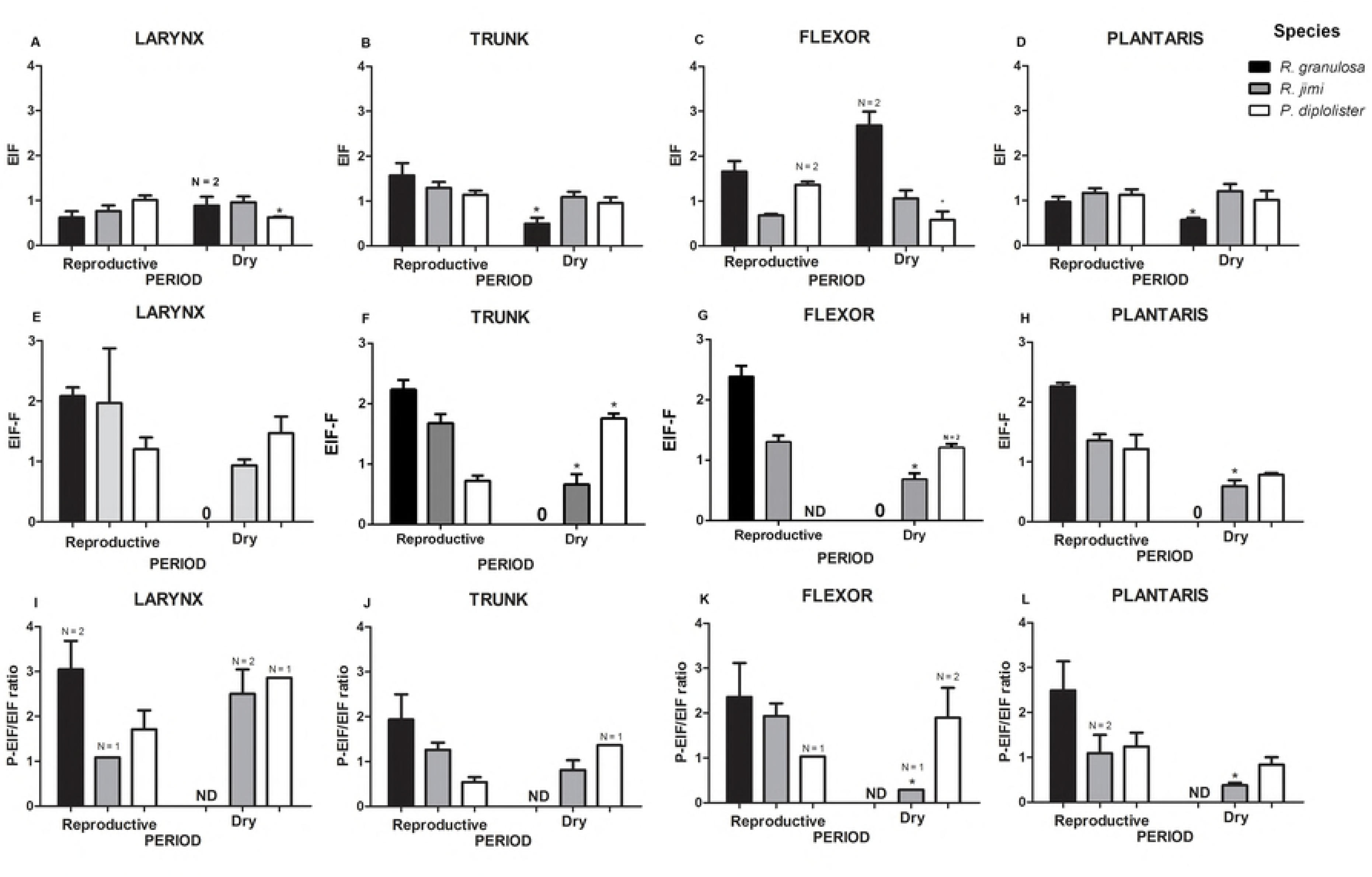
Relative intensity of eIF2α protein levels (A) larynx, (B) trunk, (C) flexor and (D) plantaris; phosphorylate eIF2α protein levels (E) larynx, (F) trunk, (G) flexor and (H) plantaris; phosphorylate eIF2α/ eIF2α ratio (I) larynx, (J) trunk, (K) flexor and (L) plantaris in *R. granulosa, R. jimi*, and *P. diplolister* during reproductive and dry period. n = 3–10, except when it is indicated. Zero indicates protein levels below the detection limit. ND - not determined because of the lack of samples. Asterisks indicate significant difference between reproductive and dry periods (P < 0.05).

Phosphorylated eukaryotic initiation factor 2-α (p-eIF2α) showed lower expression during the dry period in trunk, flexor and plantaris muscles (F_1,13_ = 0.94, P = 0.003; F_1,9_ = 0.23, P = 0.006; F_1,10_ = 0.47, P = 0.004, respectively) (Fig 2E-H), but not in the larynx from *R. jimi* (F_1,4_ = 3.52, P = 0.13). In contrast, p-eIF2α levels were elevated in the muscles of *P. diplolister* during the dry period compared to the reproductive one (Fig 2E-H), and this trend was significant in the trunk (F_1,6_ = 0.30, P = 0.003) and marginally significant in the flexor muscle (F_1,1_ = 83.38 P = 0.069) but not in the larynx (F_1,8_ = 0.673, P = 0.436) or plantaris (F_1,12_ = 0.506 P = 0.49). Notably, p-eIF2α levels were below the detection limit in all muscles from *R. granulosa* during the dry period (at 50 µg of total protein per lane) (Fig 2E-H). Comparisons between different muscle types within each species and each study season showed similar p-eIF2α expression among different muscle types in *R. jimi and R. granulosa* during the dry and reproductive periods, and in *P. diplolister* during the reproductive period (Table 1). However, during the dry period the trunk muscle of *P. diplolister* had significantly higher levels than flexor and plantaris, and levels of p-eIF2α in the plantaris tissue was below that in all other studied muscle types (Table 1, S3D Fig).

The ratio of p-eIF2α to total eIF2α levels was lower during the dry period in all muscle types of *R. granulosa* (reflecting the non-detectable levels of p-eIF2α during the dry period) and in the flexor and plantaris muscles *R. jimi* (F_1,3_ = 0.518, P = 0.02; F_1,4_ = 0.257, P = 0.034), but not P. *diplolister* (P > 0.05) (Fig 2I-L). The ratio of p-eIF2α to total eIF2α did not significantly vary among the different muscle types when compared within the same season in *P. diplolister* and *R. granulosa* (Table 1). In *R. jimi* during the dry period, larynx muscle showed higher p-eIF2α/ eIF2α ratio when compared to the other muscles (S3E Fig; Bonferroni P = 0.05); this difference was not significant during the reproductive period (Table 1).

Heat shock protein 60 (HSP60) showed lower expression during the dry period in the flexor from *R. jimi* (F_1,11_ = 1.12, P = 0.001), and in the larynx and trunk of *P. diplolister* (F_1,7_ = 0.33, P = 0.047; F_1,8_ = 0.11, P = 0.008) (Fig 3A-D). In all other studied tissue/species combinations, no significant differences in HSP60 levels were found between the reproductive and dry periods (P > 0.05). Notably, the trunk muscles of *R. jimi* and *R. granulosa* had lower HSP60 levels compared to other muscle types in the reproductive season (Table 1; S4A and C Fig). During the dry period, HSP60 levels in the trunk muscle of *R. jimi* were similar to that in the larynx and flexor while HSP60 levels in the plantaris muscle were higher than in the larynx and flexor (Table 1; Bonferroni P = 0.052) (S4B Fig). In *P. diplolister*, HSP60 were similar among different muscle types within each respective studied season (P > 0.05).

**Fig 3.**
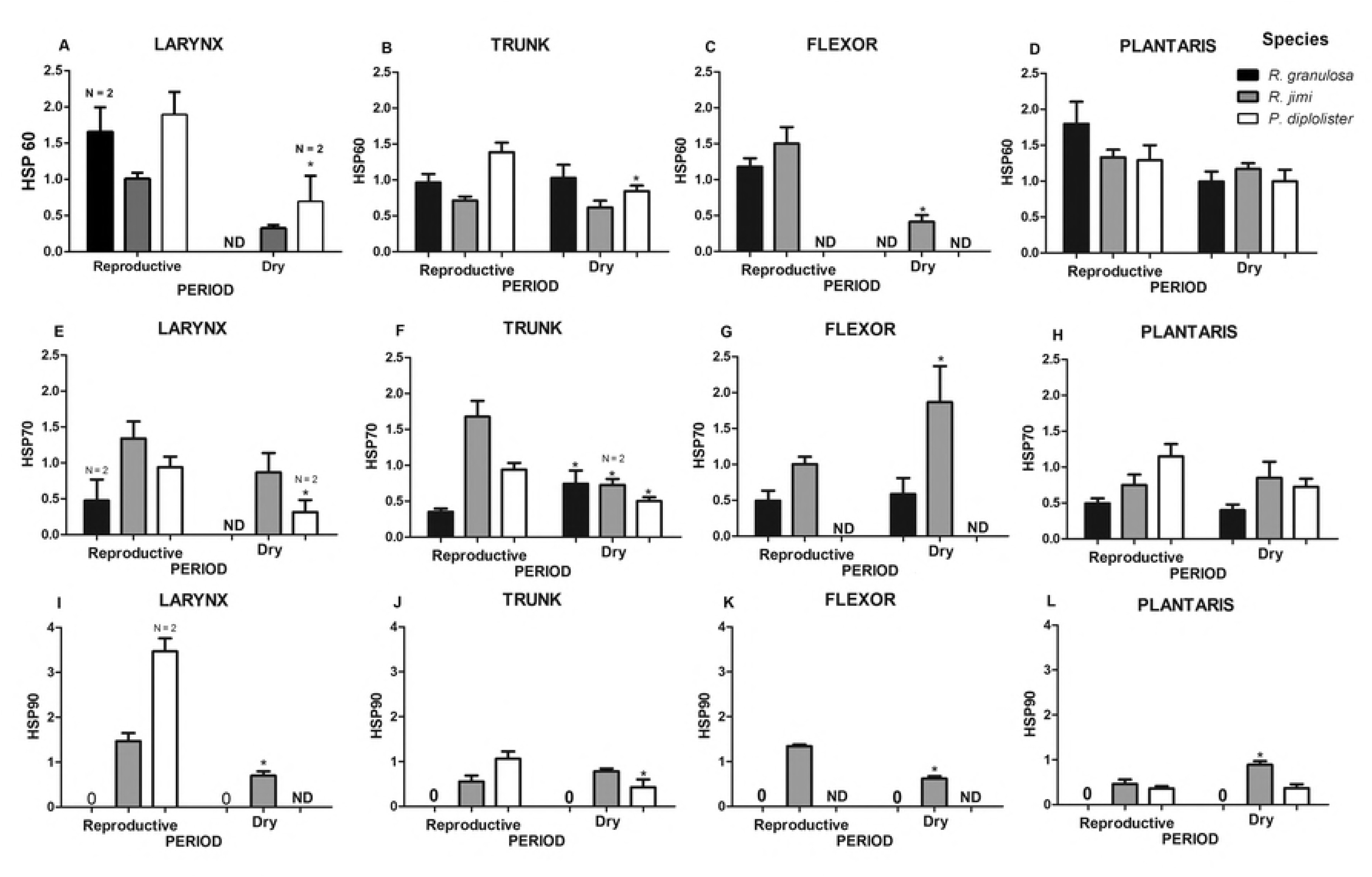
Relative intensity of HSP 60 protein levels in (A) larynx, (B) trunk, (C) flexor, (D) plantaris, HSP 70 protein levels in (E) larynx, (F) trunk, (G) flexor, (H) plantaris, HSP 90 protein levels in (I) larynx, (J) trunk, (K) flexor and (L) plantaris of *R. granulosa, R. jimi*, and *P. diplolister* during reproductive and dry periods. Data are means ± S.E.M., n = 3–11, except when indicated. ND indicates not determined because of the lack of samples. Zero indicates protein levels below the detection limit of the method employed. ND - not determined because of the lack of samples. Asterisks indicate significant difference between reproductive and dry periods (P < 0.05).

Heat shock protein 70 (HSP70) showed lower expression during the dry period compared to the reproductive season in the larynx and trunk of *P. diplolister* (F_1,9_ = 0.42, P = 0.03; F_1,13_ = 0.191, P = 0.02) (Fig 3E-H). Similarly, the trunk muscles of *R. jimi* had lower HSP70 levels during dry period compared to the reproductive one (F_1,7_ = 0.19, P = 0.02). The flexor muscle of *R. jimi* showed higher HSP70 expression during the dry period (F_1,6_ = 10.27, P = 0.019), while larynx displayed similar HSP70 expression along the year (F_1,5_ = 2.05, P = 0.21) (Fig 3E-H). *R. granulosa*‘s trunk muscles (but not the plantaris, flexor or larynx) showed higher HSP70 levels during the dry period compared to the reproductive season (F_1,8_ = 8.35, P = 0.02) (Fig 3E-H). Within each species and study season, only *R. jimi* showed differences in HSP70 levels among different muscle types during the reproductive season, with the highest levels in the trunk and the lowest levels in the plantaris muscle (Table 1; S4D Fig; Bonferroni P = 0.006).

Heat shock protein 90 (HSP 90) showed lower expression during the dry period compared with the reproductive period in the larynx and flexor muscles of *R. jimi* (F_1,8_ = 0.27, P = 0.007; F_1,11_ = 0.37, P = 0.001) and in the trunk muscles from *P. diplolister* (F_1,1_ = 1.02, P = 0.03; F_1,6_ = 0.44, P = 0.042) (Fig 3I-L). In contrast, in the plantaris muscle of *R. jimi* lower levels of HSP90 were found during the reproductive period compared with the dry one (F_1,2_ = 0.13 P = 0.045). HSP90 could not be detected in *R. granulosa* due to the lack of the cross-reactivity of the antibody. Comparison among different muscle types within the same season showed elevated HSP90 levels in the larynx and flexor compared to the trunk and plantaris of *R. jimi* during the reproductive period (Bonferroni P = 0.013), and in the larynx of *P. diplolister* compared to all other tissue types, also during the reproductive period (Bonferroni P = 0.007) (S4E and F Fig, Table 1). In all other studied species/season combinations, no significant differences in HSP90 levels were found between different muscle types (P > 0.05).

### Aerobic capacity

Activity of cytochrome c oxidase (COX), which can serve as a marker of the mitochondrial density and thus aerobic capacity of the tissue, was higher in the trunk and plantaris muscles of *R. jimi* compared to *R. granulosa* and *P. diplolister* (F_2,22_ = 7.000, P = 0.005 and F_2,43_ = 9.616, P = 0.001, for the dry and the reproductive period respectively) (Fig 4). Notably, COX activity tended to be lower in the trunk muscle than in the plantaris muscle across the species during the dry period (by 40-45% in *R. granulosa* and *P. diplolister* and by 68% in *R. jimi),* although this difference was significant only for *R. jimi* (F_1,22_ = 5.334, P = 0.03). Comparing within species, COX activity was marginally higher during reproductive season than during the dry season in the trunk *R. granulosa* (t = 2.194, df = 5, P = 0.07), but not in the plantaris (t = 0.132, df = 1, P = 0.90). *Rhinella jimi* and *P. diplolister* did not display differences in the COX activity between seasons (P > 0.05).

**Fig 4.**
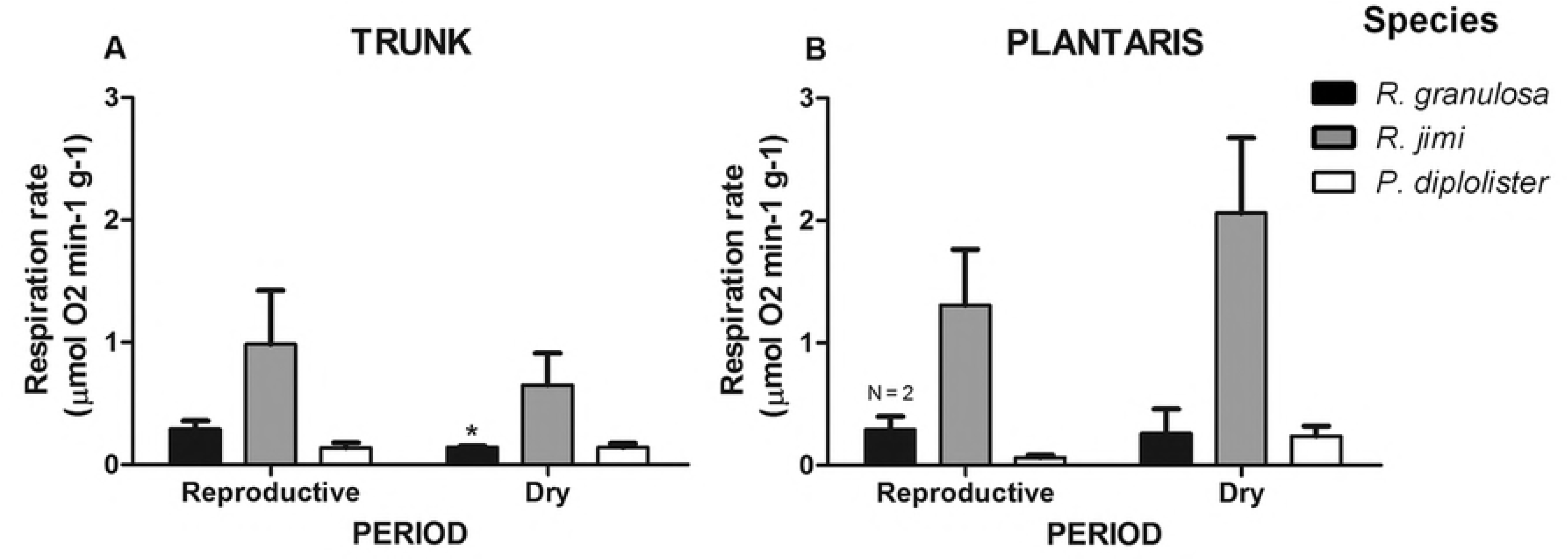
Activity of cytochrome c oxidase (COX) in trunk (A) and plantaris (B) muscles during reproductive and dry periods in *R. granulosa, R. jimi*, and *P. diplolister*. n = 3–11, except when indicated. Asterisk indicates significant difference between periods (P < 0.05).

## Discussion

Our results indicate the importance of metabolic regulators mediating the muscle maintenance and function during the drastic seasonal variation faced by the Caatinga anurans. The toads that remain active during the dry period appear to maintain muscles through more energy extensive pathways including elevated protein synthesis, while the aestivating frogs employs energy conservation strategy that involves suppression of protein synthesis, decrease in the chaperone expression and higher total expression of AMPK. These adjustments are consistent with their lower metabolic rates [12] and need for saving energy during aestivation. All three studied species activate cell survival pathways during the dry period in the muscles likely to prevent muscle atrophy. All three studied species thereby maintain the muscle capacity throughout the year, despite the resource limitation. These strategies are important considering the unpredictability of the reproductive event and the need to rapidly engage the muscular activity in response to the rain event triggering reproduction.

### Cellular survival and protein synthesis pathways in the muscle

The protein kinase B (AKT) is an important signaling protein involved in cellular survival pathways in the muscle [28]. AKT expression was elevated during the dry period in the trunk and plantaris muscles from all studied species and in the larynx in the aestivating species, but not in flexor. The most pronounced increases of AKT was in the aestivating species, *P. diplolister*. Possibly, activation of AKT in the flexor may occur later as the dry season progresses (similar to the delayed activation of the AKT pathway in some skeletal muscle types of a hibernating mammal *Marmota flaviventris* [29]; this hypothesis remains to be tested with regard to the Caatinga anurans. Similarly to our present results, elevated expression of AKT was reported in the foot muscle and hepatopancreas of estivating snails [30] and in the liver of a frog *Rana sylvatica* exposed to freezing [31]. AKT promotes cell survival, downregulates pro-apoptotic factors [32,30,33,31,34], and activates the cascade involved in cell cycle arrest and quiescence [35]. Therefore, the upregulation of AKT in the Caatinga anurans during the dry period could be important for preventing the muscle atrophy when resources are limited. Notably, the upregulation of AKT during the dry period goes hand-in-hand with an increase of the phosphorylated form of eIF2α (p-eIF2α) in the trunk muscle and/or a decrease of the total eIF2α in the larynx and flexor muscles of the estivating species, *P. diplolister*. This agrees with the earlier findings showing that phosphorylated eIF2α can facilitate AKT activation thereby promoting cell survival [36]. Furthermore, the eIF2α is an essential initiation factor in protein synthesis which controls the translation rates and becomes inactivated by phosphorylation [37]. Thus, low levels of eIF2α and/or elevated expression of p-eIF2α indicate suppression of the protein synthesis in the muscles of *P. diplolister* during aestivation. The stable level of eIF2α and p-eIF2α in the other studied *P. diplolister* muscles might indicate that the regulation of the protein synthesis in these tissues might be dependent on alternative mechanisms such as control of the translation elongation [38] or ribosome (dis)assembly [39]. Suppression of the protein synthesis is a common energy-saving mechanism in estivating species [38] and has been observed in desert amphibians *Neobatrachus centralis* [40] and in estivating snails *Otala lactea* [41,42]. Earlier studies on estivating frogs (*Cyclorana alboguttata)* showed that muscles are protected against atrophy during prolonged (9 months) estivation with no decline in muscle mass, cross-sectional area or fiber number [7]. Our study in *P. diplolister* suggests a possible mechanism for this protection involving the coordinated suppression of the protein synthesis to conserve energy reserves and activation of the cell survival pathways to prevent loss of the muscle cells.

The protein synthesis was activated during the dry period in the muscles of *R. jimi* and *R. granulosa* (two species that remain active throughout the year) as indicated by a decline in the amount of inactive p-eIF2α in all studied muscle types, but not in the aestivating species *P. diplolister*. This was especially notable in *R. granulosa* where p-eIF2α levels were below the detection limits of immunoblotting during the dry period. The maintenance of foraging activity of *Rhinella* species during dry period might allow higher protein synthesis and activation of the cell survival pathways, potentially helping to build the muscle mass in preparation for the reproductive period in the two toad species. The maintenance of the muscle mass during the dry period is important for the Caatinga anurans, which start reproduction immediately after fairly unpredictable rainfall events [13]. The reproductive behavior involves strenuous calling activity (which engages the trunk muscles) in all three studied species [12]. In *P. diplolister*, males must also energetically beat legs to build foam nests for eggs deposition [12]. Despite some variation in the AKT, eIF2α and p-eIF2α levels among the muscle types, the seasonal patterns of expression of these proteins were generally consistent in different muscles within each studied species. These results indicate that the molecular mechanisms of the muscle maintenance during the resource-limited dry season are similar in the locomotor muscles (i.e. plantaris) and the reproductively-related muscles (such as trunk, flexor and larynx).

### Indices of energy status

Elevated expression of AMPK in muscle tissues (indicative of the cellular energy stress) was observed during the dry period in all muscles of *R. jimi* and in plantaris muscle of *P. diplolister*. An increase in AMPK levels is common during the resource- and energy-limited periods in many organisms including hibernating mammals [43,44] and frogs exposed to hypothermia, hypoxia, freezing, dehydration or anoxia [45,46,44]. Furthermore, AMPK suppresses energy-demanding metabolic processes [47,22] and induces cell cycle arrest [12,22,44], which also contributes to energy savings. The reasons for the differences in AMPK response between the two non-estivating active species are not known. Furthermore, the expression of phosphorylated and total AMPK might not always be in the same direction which limits our conclusions on the potential changes in the active form of this protein (since the phosphorylated form of AMPK could not be measured in our present study due to the lack of antibody cross-reactivity with the anuran protein). Different responses regarding AMPK activation in anuran species submitted to the same freezing condition has been reported in *Rana perezi* and *Rana sylvatica* [45,46]. Additionally, during reproductive period, AMPK expression was particularly elevated in trunk of *P. diplolister* compared to other muscle types, which might indicate a metabolic stress due to high energy demand of the trunk muscles due to calling activity [48,49,50]. However, this hypothesis remains to be tested.

COX activity (indicative of the mitochondrial density) was considerably higher in the muscles of the largest of the three studied species (*R. jimi)* compared to *P. diplolister* or *R. granulosa*. COX activity was especially high in the plantaris muscle of *R. jimi,* which is consistent with higher locomotor capacity of large toads from *Rhinella marina* group of species [51]. Generally, the mitochondrial COX capacity of the frogs’ muscles was maintained at the same level in the reproductive and dry period except for a small but significant decline in the COX activity in the trunk muscle of *R. granulosa* during the dry period. This indicates that the aerobic capacity of the locomotor as well as the reproductive muscles is maintained throughout the year despite the energy and resource limitation in the dry season.

### Expression of molecular chaperones

Molecular chaperones involved in the folding of nascent and damaged proteins (including a mitochondrial HSP60 and cytosolic HSP70 and HSP90) were expressed at lower levels in the muscles of the estivating *P. diplolister* during the dry period. A decrease in HSP expression in aestivating frogs goes hand-in-hand with the suppressed protein synthesis in the muscles and may reflect lower protein turnover rates during aestivation [41,52]. Similarly, other aestivating species such as land snails also decrease HSP expression during quiescence [53, 54] and sharply increase it during arousal [55].

The higher expression of HSPs in the muscles trunk of *P. diplolister* during the reproductive period might be attributed to the stress caused by the exercise from calling behavior and high steroid receptor expression levels [56,57,58]. Calling is a highly energetic demanding aerobic exercise for anuran males [59,48], and calling effort is positively correlated with plasma levels of androgens and corticosterone [60, 65, 66]. Furthermore, anurans show increased expression of steroid hormones during reproductive season [65,67], which are associated with HSP in the inactive state [68,69,70]. Thus, elevated expression of HSPs during the reproductive season in *P. diplolister* might also reflect the high levels of steroids during this period. However, the pattern of HSP levels in the muscles of *R. jimi* or *R. granulosa* males are less clear, considering they also display high steroid levels during the reproductive season. Overall, HSP levels of in the muscle tissues in the two anurans species that are year-round active showed relatively little variation and no consistent pattern of seasonal change between different muscle types. This indicates the lack of strong unfolded protein response and thus maintenance of the protein homeostasis during the period of reproductive activity as well as during times of resource limitation in these species.

## Conclusions and perspectives

The differential responses of the cell signaling and stress response pathways found in the muscles of the three studied species of desert anurans reflect differences in their life habit and activity levels as well as the common need to maintain the muscle capacity in an extremely seasonal environment of the Caatinga. Activation of the cell survival pathways is the most consistent response in the muscles of all three studied species during the dry period likely playing a role in preventing muscle atrophy during the resource limitation. Expression of the regulators of protein homeostasis (including chaperones and regulators of the protein synthesis) reflect different levels of resource limitation, so that the less resource-limited active species upregulate protein synthesis and maintain high levels of chaperones during the dry periods in the muscle while a severely resource-limited aestivating species shuts down the protein synthesis to conserve energy. Contrary to our prediction, we did not observe differential metabolic regulation or trade-off between the reproductive and locomotor muscles. Future studies are needed to determine whether the muscle maintenance during the dry period is prioritized over that of other tissues and whether potential trade-offs exist between the support of the muscle capacity throughout the year and other fitness-related functions in desert frogs.

## Acknowledgements

We are grateful to Assis, F. and Cassettari, B.O. for support in the fieldwork; and Ivanina, A.V. and Martins, A.N. for sharing their time, knowledge and expertise.

## Author contributions

I.S., C.B.M and F.R.G. conceived the study, designed the experiments and contributed substantially to interpreting the data. C.B.M and I.S. collected the data; C.B.M. analyzed the data; C.B.M, I.S. and F.R.G. wrote the manuscript, and take full responsibility for the content of the paper.

## Supporting information

**Fig S1. AMPK expression during (A) reproductive and (B) dry period for *R. jimi*, AKT expression during (C) reproductive period for *R. jimi*; Total eIF2α expression during (D) reproductive and (E) dry period for *P. diplolister;* Phosphorylated eIF2α expression during (F) reproductive and (G) dry period for *R. jimi*; HSP60 expression during (H) reproductive and (I) dry period for *R. jimi*; HSP70 expression during (J) reproductive period for *R. jimi*; and HSP90 expression during (K) reproductive period for *R. jimi*.**

**Fig S2. Relative intensity of AMPK protein levels among different muscles in *P. diplolister* during reproductive period. Data are means ± S.E.M., n = 7–10. ND indicates not determined because of the lack of samples.** Different letters indicates significant difference (P < 0.05).

**Fig S3. Relative intensity of eIF2α protein levels among different muscles in *R. granulosa* during (A) reproductive and (B) dry period and *R. jimi* during (C) reproductive period; phosphorylate eIF2α among different muscles in *P. diplolister* during (D) dry period; and phosphorylate eIF2α/eIF2α ratio among different muscles in *R. jimi* during (E) reproductive period.** Data are means ± S.E.M., n = 3– 10, except when otherwise indicated. Asterisks and different letters indicates significant difference (P < 0.05).

**Fig S4. Relative intensity of HSP 60 protein levels among different muscles in *R. jimi* during (A) reproductive and (B) dry period and in *R. granulosa* during (C) reproductive period; HSP 70 protein levels in *R. jimi* during (D) reproductive period; and HSP 90 in *R. jimi* during (E) reproductive period and in *P. diplolister* during (F) reproductive period.** Data are means ± S.E.M., n = 4–8, except when otherwise indicated. ND indicates not determined because of the lack of samples. Asterisks and different letters indicates significant difference (P < 0.05).

